# Expression of coffee florigen *CaFT1* reveals a sustained floral induction window associated with asynchronous flowering in tropical perennials

**DOI:** 10.1101/2021.11.04.466514

**Authors:** Carlos Henrique Cardon, Raphael Ricon de Oliveira, Victoria Lesy, Thales Henrique Cherubino Ribeiro, Catherine Fust, Luísa Peloso Pereira, Joseph Colasanti, Antonio Chalfun-Junior

## Abstract

The behavior of florigen(s) and environment-influenced regulatory pathways that control flowering in tropical perennials with complex phenological cycles is poorly understood. Understanding the mechanisms underlying this process is important for food production in the face of climate change. To explore this, homologs of *Arabidopsis* florigen *FLOWERING LOCUS T* (*CaFT1*) and environment-related regulators *CONSTANS* (*CO*)*, PHYTOCHROME INTERACTING FACTOR 4* (*PIF4*) and *FLOWERING LOCUS C* (*FLC*) were isolated from *Coffea* sp. L. (Rubiaceae). Overexpression of *CaFT1* in *Arabidopsis* showed typical early-flowering and yeast two hybrid studies indicated CaFT1 binding to bZIP floral regulator, FD, demonstrates that CaFT1 is a coffee orthologue of florigen. Expression of *CaFT1* and floral regulators were evaluated over one year using three contrasting genotypes: two *C. arabica* and one *C. canephora*. All genotypes showed active *CaFT1* transcription from February until October, indicating a potential window for floral induction. *CaCO* expression, as expected, varied over the day period and monthly with day length, whereas expression of temperature-responsive homologs, *CaFLC* and *CaPIF4*, did not correlate with temperature changes. Using coffee as a model, we suggest a continuum of floral induction that allows different starting points for floral activation, which explains developmental asynchronicity and prolonged anthesis events in tropical perennial species.

**Highlight:** Coffee florigen CaFT1 and related regulators revealed an extended floral induction window clarifying the asynchronicity and influence of environment for flowering in tropical perennial crops, providing perspectives to its control.

## INTRODUCTION

In flowering plants, the transition of vegetative meristems (VMs) to a reproductive state is mainly controlled by members of the phosphatidylethanolamine-binding protein (PEBP) family that act as transcriptional coactivators (Cao *et al.*, 2016). The *Arabidopsis FLOWERING LOCUS T* (*FT*) gene is the first PEBP member identified to encode a florigen; that is, FT translocation via phloem from source leaves to VMs induces floral meristem (FM) production (Wigge *et al.*, 2005; Corbesier *et al.*, 2007). To ensure proper reproductive timing, the FT pathway integrates inductive exogenous stimuli, such as photoperiod, temperature, gibberellins (GA) and vernalization, together the endogenous autonomous pathway (Amasino, 2010; Song *et al.*, 2008; Kim *et al.*, 2009; Andrés and Coupland, 2012). The flowering process is very well explored in model plants such as *Arabidopsis*, however, it is poorly understood in perennial plants that have more complex reproductive cycles, such as *Coffea* sp. (Rubiaceae).

An example of this complexity is that perennial plants must maintain simultaneously in a branch the shoot apical meristem (SAM) in a vegetative state to guarantee the continuity of plant vegetative growth and reproduction in subsequent years, whereas older meristems at the base of branches form FMs and then flourish (Amasino, 2009; Albani and Coupland, 2010). Thus, this mode of reproduction is intrinsically connected with plant architecture indicating a precise molecular control to determine the different forms of meristems (Wang *et al.*, 2009; Albani and Coupland, 2010). The coffee phenological cycle is biennial, which means that it takes two years to produce fruits (Teketay, 1999; Camargo and Camargo, 2001). The first year is characterized by the floral induction and floral development until anthesis (or bloom) which, in *Brazil*, occurs from January to March and in September, respectively (Camargo and Camargo, 2001; Majerowicz and Söndahl, 2005; Morais *et al.*, 2008). Importantly, the floral induction period is based on morphological observations, when vegetative and floral buds are distinguishable (de Oliveira *et al.*, 2014), but without precise molecular characterization of the inductive floral pathway (reviewed by López *et al.*, 2021 in press).

Organogenesis during coffee floral development was previously characterized (Majerowicz and Söndahl, 2005; de Oliveira *et al.*, 2014) and shown to be affected by environmental conditions that result in uneven development and sequential flowering (Camargo and Camargo, 2001; DaMatta and Ramalho, 2006; DaMatta *et al.*, 2007). For example, in the G4 stage coffee buds enter a latent state during the cold and dry period of Brazilian winter in June to August (Wormer and Gituanja, 1970; Camargo and Camargo, 2001; Morais *et al.*, 2008). After this period, bud growth resumes in a quick passage from the G4 to G6 following the anthesis that coincides with the rain period around September. Thus, this suggests that coffee flowering is controlled by different factors, such as low temperatures and plant water potential (Crisosto *et al.*, 1992; Majerowicz and Söndahl, 2005; DaMatta and Ramalho, 2006), in addition to photoperiod, as commonly observed in angiosperms. Nevertheless, from the molecular point of view, the mechanism involved in the perception of these environmental stimuli and its relationship with the *FT-*dependent inductive pathway has never been explored in coffee.

The PEBP family of proteins can be divided into five subgroups defined by *Arabidopsis* family members: *FLOWERING LOCUS T* (*FT*)*, TWIN SISTER OF FT* (*TSF*)*, TERMINAL FLOWER 1* (*TFL1*)*, BROTHER OF FT AND TFL* (*BFT*) and *MOTHER OF FT AND TFL* (*MFT*). These phosphatidyl ethanolamine-binding protein (PEBP) genes have important roles in plant development, most notably floral induction and inflorescence architecture (Karlgren *et al.*, 2011; Zhu *et al.*, 2021). Accordingly, PEBP members are highly conserved between flowering species, especially *FT* and *TFL1* sequences (Turck *et al.*, 2008; Wickland and Hanzawa, 2015). Interestingly, FT and TFL1 proteins are highly similar, yet they have opposite functions. That is, FT promotes floral meristem formation whereas TFL1 represses flowering and maintains the inflorescence in an indeterminate state (Kaneko-Suzuki *et al.*, 2018; Nakamura *et al.*, 2019). These antagonistic roles are due FT and TFL1 competition for the bZIP transcription factor FD and 14-3-3 proteins to create a floral activation complex (FAC) or a floral repression complex (FRC), respectively (McGarry and Ayre, 2012; Kaneko-Suzuki *et al.*, 2018). Another difference is that FT protein accumulates in leaves and then moves long distances to the SAM (Yoo *et al.*, 2013), whereas *TFL1* is expressed in a subdomain of the SAM (Bradley *et al.*, 1997).

In *Arabidopsis*, long-day (LD) induction of *FT* expression is largely dependent on the B-box transcription factor CONSTANS (CO), which is modulated by the circadian clock and day length (Amasino, 2010; Andrés and Coupland, 2012; Liu *et al.*, 2014). Under LD conditions, CO protein accumulates to a threshold sufficient to activate *FT* expression, whereas in short days (SD) CO is degraded via the 26S proteasome complex and does not activate *FT* expression (Suárez-López *et al.*, 2001; Valverde *et al.*, 2004; Zuo *et al.*, 2011). Other key transcription factors (TFs) regulating *FT* are PIF4 and FLC, both responsive to temperatures changes and activity depending on epigenetic changes in chromatin (Helliwell *et al.*, 2006; Kumar *et al.*, 2012; Madrid *et al.*, 2021). PIF4 is responsive to warmer temperatures and positively regulates *FT* expression, whereas FLC is a repressor that is inactivated by prolonged cold periods (vernalization) allowing *FT* expression and flowering initiation (Michaels *et al.*, 2005; Amasino, 2010). *FLC* homologs together with other MADS-box genes were described in *C. arabica* (de Oliveira *et al.*, 2010; 2014) and related to drought (Barreto *et al.*, 2012) that occurs in the cold period of the Brazilian winter. However, the interplay between the FT pathway and environmental signals through the identification and analysis of *CO*, *PIF4* and *FLC* and their roles in modulating floral induction is poorly understood in perennials and has never been explored in coffee.

In addition to environmental effects, another regulatory factor is sugar signaling, which plays an important role in plant development, including flowering regulation (Corbesier *et al.*, 1998; Lastdrager *et al.*, 2014; Sheen, 2014). Day length directly affects sucrose and starch accumulation as a consequence of photosynthesis, which has been correlated with reproductive induction (Corbesier *et al.*, 1998). Accordingly, varying day length of different seasons is directly associated with the transition from vegetative to reproductive time in plants (reviewed by Andrés and Coupland, 2012). A possible role for sucrose as a signal for flowering is implicated by its influence on *FT* expression (Moghaddam and Van den Ende, 2013). Sucrose has been shown to act in leaves, downstream of *CO* and upstream to *FT,* to affect flowering (Corbesier *et al.*, 1998; Ohto *et al.*, 2001). Moreover, transcriptomic studies of maize floral regulator *indeterminate1* (*id1*) shows a correlation between sugar and starch metabolism and floral induction in this autonomously flowering plant (Coneva *et al.*, 2007, 2012; Minow *et al.*, 2018).

Florigen and *FT* orthologs in diverse plant species are key integrators of reproductive meristem specification via interconnecting molecular pathways that determine the proper floral induction window (Amasino, 2009; He *et al.*, 2020). Despite conservation across plant species, variable expression patterns of *FT* homologs have been reported (Coelho *et al.*, 2014; Cao *et al.*, 2016; Wolabu *et al.*, 2016; Štorchová *et al.*, 2019). This variable expression likely reflects the variety of flowering evolutionary strategies required to tailor flowering time and floral architecture diversification to particular geographical locations (Pin and Nilsson, 2012; Wickland and Hanzawa, 2015; Jin *et al.*, 2020). Very little research on the mechanisms underlying floral induction and its interplay with environmental-dependent pathways has been carried out in *Coffea* sp, a tropical perennial crop of considerable socio-economic importance (IOC, 2021).

Here we report the first characterization of homologs of the key floral regulators *FT*, *CO*, *PIF4,* and *FLC* from *C. arabica*, *C. canephora* and *C. eugenioides.* Through transgenic analysis and protein interaction assays we demonstrate that *CaFT1* from coffee is a functional floral inducer (an *FT* ortholog). To determine the precise floral induction window and establish a possible correlation with environmental signals, expression of *CaFT1* was determined during a year at two daily time points together with *FT* regulators *CaCO*, *CaPIF4,* and *CaFLC*. Expression was analysed in three coffee genotypes with contrasting flowering patterns, two *C. arabica* cvs. Acauã and IAPAR59 and *C. canephora* cv. Conilon, to examine related intra- and interspecific variation. In addition, to investigate a possible connection between sugar levels, gene expression and floral development, carbohydrate content was determined at the same time points for all genotypes. Based on these results we propose that an extended floral induction window for coffee could explain the asynchronous development and sequential flowering in this perennial species. Whereas *CaFT1* expression was correlated to short day photoperiod and cold (Brazilian winter), a similar pattern was not detected in the expression profiles of *CaCO*, *CaPIF4,* and *CaFLC,* indicating a more complex process regulating the flowering process. Thus, this work contributes to our understanding of the floral transition in perennial species and could support future strategies aimed at mitigating asynchronous flowering.

## MATERIAL AND METHODS

### Plant material

Plant material used for RNA extraction, cloning, gene expression, carbohydrate content (total soluble sugar, sucrose, reducer sugar, and starch) was obtained from 4-year-old coffee plants of three different cultivars established at the National Institute of Science and Technology (INCT-Cafe) experimental field at the Federal University of Lavras, Brazil (21°23’S, 44°97’W): *C. arabica* genotypes IAPAR 59 and Acauã, and *C. canephora* Conilon. All plants were cultivated under nutritional and pest control conditions recommended for coffee (Vieira, 2008). Three biological repetitions of each coffee cultivar were distributed randomly in the field. Each sampling consisted of a mix of three completely expanded leaves collected and immediately immersed in liquid nitrogen and stored at −80 °C until RNA extraction. Samples were collected at five-time points (December 2016, February, April, June, and October 2017) over a two-day period, at 6:00 am and 5:00 pm, considered the start and end of the day, totaling 90 samples. *Arabidopsis thaliana* var. Landsberg *erecta* (L*er*) was used for *CaFT1* heterologous expression studies following Coelho *et al.* (2014) plant growth conditions. Plants were grown in growth chambers under 16 hrs light / 8 hrs dark at 22 °C and 60 % humidity.

### *In silico* and phylogenetic analysis

Searches for coffee homologs of *FLOWERING LOCUS T* (*FT*), *CONSTANS* (*CO*), and *PHYTOCHROME INTERACTING FACTOR 4 (PIF4)* were performed by sequence comparisons using the BLAST tool (Johnson *et al.*, 2008). First, the described genes *FT* (At1G65480), *CO* (At5G15840), and *PIF4* (At2G43010), and *FLC* (At5g10140) from *A. thaliana* were used as queries (Lamesch *et al.*, 2012) against the *C. arabica* (NCBI:txid13443) annotated genome deposited at the National Center for Biotechnology Information (NCBI) and the Coffee Genome Hub (Denoeud *et al.*, 2014). To enrich searches for new coffee sequences phylogenetic analysis also was performed with different homologs from various species, including *Arabidopsis thaliana, Solanum lycopersicum, Brassica napus, Jatropha curcas, Nicotiana tabacum, Glycine max, Oryza sativa, Zea mays, Sorghum bicolor, Solanum tuberosum* and *Populus nigra*. The sequence for coffee *FLC*, *CaFLC*, was retrieved from the previously reported sequence (de Oliveira *et al.*, 2010).

Putative homologous sequences (>80% similarity and e-Value < 0,005) were aligned by ClustalX2 (Larkin *et al.*, 2007), analyzed by Genedoc software (Nicholas and Nicholas, 1997), and with the translated nucleotide sequence to protein, phylogeny was inferred with the nearest neighbour joining method in MEGA-X (Kumar *et al.*, 2018). Duplicate sequences were deleted from the first phylogenetic tree and only considered *A. thaliana* and *S. lycopersicum* sequences for the final tree (Fig. S1). The IDs of the used sequences are presented in the legend of Fig. S1. To provide further evidence of the putative coffee FT as a floral inducer, key amino acids were identified in alignment with CETS members from *A. thaliana* and *S. lycopersicum* (Fig. S2). The conserved amino acids that distinguish the related proteins FT and TFL1, a floral inducer and a repressor respectively (Ahn *et al.*, 2006; Wickland and Hanzawa, 2015; Jung *et al.*, 2016), are shown in Fig. S2. Similar analyses were made to show the conserved amino acids between coffee proteins and homologs of CO and PIF4 (Fig. S3 and S4).

### Gene cloning

Total RNA from coffee leaves was isolated using Concert™ Plant RNA Reagent (Invitrogen) following the manufacture’s recommendation. RNA concentration and purity were measured by spectrophotometric analysis (*GE NanoVue*™ Spectrophotometer). All samples were treated with Ambion DNase I (RNase-free) kit (Thermo Fisher) and cDNA synthesized using High-Capacity cDNA Reverse Transcriptase Kit (Thermo Fisher) following the manufacturer’s recommendation. Primers were designed for gene isolation with the putative *CaFT1* sequence previously identified by *in silico* analysis and including the restriction sites *Spe*I in the forward primer and *BsrG*I in the reverse (5’Fw = ATGCCTAGAGGGGGAGGAGA; 5’Rv = TTATCGTCTTCTGCCTC). The Polymerase Chain Reaction (PCR) was carried out using *iProof* High-Fidelity *DNA Polymerase* (Bio-Rad) following the manufacture’s protocol. PCR fragments were isolated from 1 % agarose gel after electrophoresis and purified by *GeneJET Gel Extraction Kit* (Thermo Fisher). The fragment generated was inserted into the PJET1.2/blunt cloning vector (Thermo Fisher) and transferred to pK2WG7 plasmid (Thermo Fisher), both previously digested using restriction enzymes, and then ligated with T4 DNA ligase (New England Biolabs – NEB). All procedures followed the manufacturer’s instructions.

### Plant transformation

Arabidopsis var. Landsberg *erecta* (L*er*) wild type plants (WT) and *ft* loss-of-function mutants were transformed to overexpress *CaFT1* using the *Agrobacterium tumefaciens* strain GV3101::pMP90 and floral dip protocol (Clough and Bent, 1998), using the isolated and cloned *CaFT1* fragment under the control of the CaMV 35S promoter (Gene cloning above section). Seeds from different transformation events were harvested separately for each background genotype. Positive transformed seeds were screened in selective growth media with Kanamycin 30 ug/mL and, after 2 days of incubation in darkness and at 4 °C, they were maintained in a growth chamber under continuous light for one week. Twenty positive T1 plants were transferred to soil and after two weeks, DNA was extracted from leaves and PCR analysis was conducted to confirm insertions. Overexpression of *CaFT1* driven by 35S CaMV was analyzed in 9 independent lines from 24 in T2 both L*er* wild type (L*er*-WT) plants as well as *ft* mutants.

### Yeast two Hybrid assay

Protein-protein interaction analysis by Yeast Two Hybrid assay with CaFT against AtFD and At14-3-3 was performed with the Matchmaker Gold Yeast Two-Hybrid System (Chien *et al.*, 1991). The *CaFT1* sequence was amplified by *iProof* High-Fidelity *DNA Polymerase* (Bio-Rad) with restriction enzyme site tag insertion (Fw-*EcoR*I and Rv-*BamH*I), inserted to pBridge plasmid by T4 DNA ligase and transformed in DH5α *E. coli* competent cells. Gold Yeast was transformed with *CaFT1*-pBridge (Bait) and *AtFD-*pGADT7-RecAB (Prey); and *CaFT1*-pBridge (Bait) and *At14-3-3-*pGADT7-RecAB (Pry). AtFD and At14-3-3 were screened from Arabidopsis GoldY2H library.

### Gene expression analysis (RT-qPCR)

The primers for Reverse Transcription quantitative Polymerase Chain Reaction (RT-qPCR), shown in Table S1, were designed using as template the *CaFT1*, *CaCO* and *CaPIF4* sequences identified in this work by *in-silico* analysis, whereas for *CaFLC*, the sequence was previously characterized by de Oliveira *et al.* (2010). Primers were designed following specifications for standard qPCR such as amplicon length (between 80 and 150 bases), having no sequences at the conserved domain, 40% to 60% GC content, and others as suggested by MIQE (Bustin *et al.*, 2009) and examined for hairpins and dimers with the Oligoanalyzer Tool IDT (https://www.idtdna.com/pages/tools/oligoanalyzer). The genes *CaUBQ2* and *CaMDH* and their described primers were used as reference genes (Martins *et al.*, 2017). RNA extraction, DNAse treatment, and reverse transcription reaction were performed as described above (Gene cloning session). RT-qPCR analysis was performed using 15 ng of cDNA to a final volume of 15 μL reaction with Rotor-Gene SYBR® Green PCR Kit (Qiagen), in a Rotor Gene-Q (R) thermocycler (Venlo, Netherlands). Mix reagents: 7.5 μL of SYBR-green (QuantiFast SYBR Green PCR Kit - Qiagen), 3.0 μL of forward and reverse gene-specific primers, 1.5 μL of cDNA at 10 ng/μL, and 3 μL of RNase-DNase-free water, resulting in at 15 μL final volume. Three biological repetitions were used, run in duplicate, and amplification was performed following manufacturer’s instructions. Relative expression differences were calculated by log2 of fold change and statistical analysis by Linear Mixed Model as described by Steibel *et al.* (2009).

### Carbohydrate Content Analysis

Carbohydrate quantification was carried out following methods adapted by Meireles da Silva *et al.*, (2014) with modifications: 100 mg of fresh frozen tissue previously powdered with liquid nitrogen were homogenized in 5 mL potassium buffer (100 mM, pH 7.0) followed by 30 min incubation in a water bath at 40 °C. Supernatant was collected after 10 min centrifugation at 10,000 xG and stored at −20 °C. Pellets were used for starch extraction by 5 mL potassium acetate buffer (200 mM, pH 4.8) resuspended with 16 units of amyloglucosidase enzyme, followed by two hours at 40 °C and 20 min of centrifuge at 10,000 xG for supernatant separation and starch quantification. Starch, sucrose, and total soluble sugars were quantified according to Dische (1962) and, reducing sugar levels according to Miller (1959).

### Statistical analysis

A Linear Mixed Model was used for statistical analysis of biochemistry quantification by the lme4 R package (Bates *et al.*, 2015), and adjustment parameters following Oliveira *et al.*, (2020) statistical methods. RT-qPCR statistical analysis was carried out by log2 of fold change by Linear Mixed Model as described by Steibel *et al.* (2009).

## RESULTS

### Identification of *FT*, *CO* and *PIF4* homologs from *Coffea* sp

To identify *FT* sequences and other genes related to floral regulatory pathways, such as *CO* and *PIF4*, we searched for homologous genes in *C. arabica* (NCBI:txid13443) and *C. canephora* (https://coffee-genome.org/) genome databases. Through phylogenetic analysis, sequences were compared to their homologs in other species to identify the most similar counterparts as putative orthologs (**Fig. S1**). Genome analysis of homologous sequences identified seven *FT-*like genes, four *CO-*like genes and four *PIF4-*like genes. The PEBP family members, which include FT-like (**Fig. S1-A**), were separated into three subgroups based on previous reports: *FT-LIKE* (*FT* and *TSF*), *TFL1-like* (*TFL1* and *BFT*) and *MFT-like* (*MFT*) (Karlgren *et al.*, 2011; Nasim *et al.*, 2017).

Sequence alignments of PEBP homologs, including those from *Arabidopsis thaliana* and *Solanum lycopersicum* (**Fig. S2**), revealed conserved amino acids in *Coffea* sequences related to FT-like floral inducers. For example, the conserved Y85 amino acid residue reported to be a hallmark of FT orthologs supports the identified coffee genes as a possible florigen (Hanzawa *et al.*, 2005). These FT-like sequences used in further functional characterization were identical in *C. arabica* and *C. canephora* and here both are referred to as *CaFT1*. Similar genomic analysis was undertaken for CO-like (**Fig. S3**) and PIF4-like genes **(Fig. S4)**, which showed conserved motifs indicating they are putative *Coffea* sp. orthologs.

### Overexpression of *CaFT1* in *Arabidopsis* and protein interaction analysis suggest that it is a *Coffea* sp. *FT* ortholog

We explored *CaFT1* function by over-expressing this gene in *Arabidopsis thaliana* Landsberg *erecta* (L*er*) to examine the effects on flowering. In addition, yeast two hybrid (Y2H) assays were performed to test conserved protein-protein interactions between CaFT1 and components of the Arabidopsis floral regulatory complexes. In 24 independent lines from T2 generations of *CaFT1* overexpression driven by the 35S CaMV promoter, nine independent lines caused a strong early flowering phenotype in both L*er* wild type (L*er*-WT) plants as well as *ft* mutants. Transgenic plants flowered 12 days after germination (DAG) in contrast to L*er*-WT plants, which flowered at 28 days (**Fig. 1A**). Accelerated transition from vegetative to reproductive development was also indicated by the reduced number of rosette leaves initiated in both L*er*-WT and *ft* mutant backgrounds (**Fig. 1B**).

**Figure 1:**
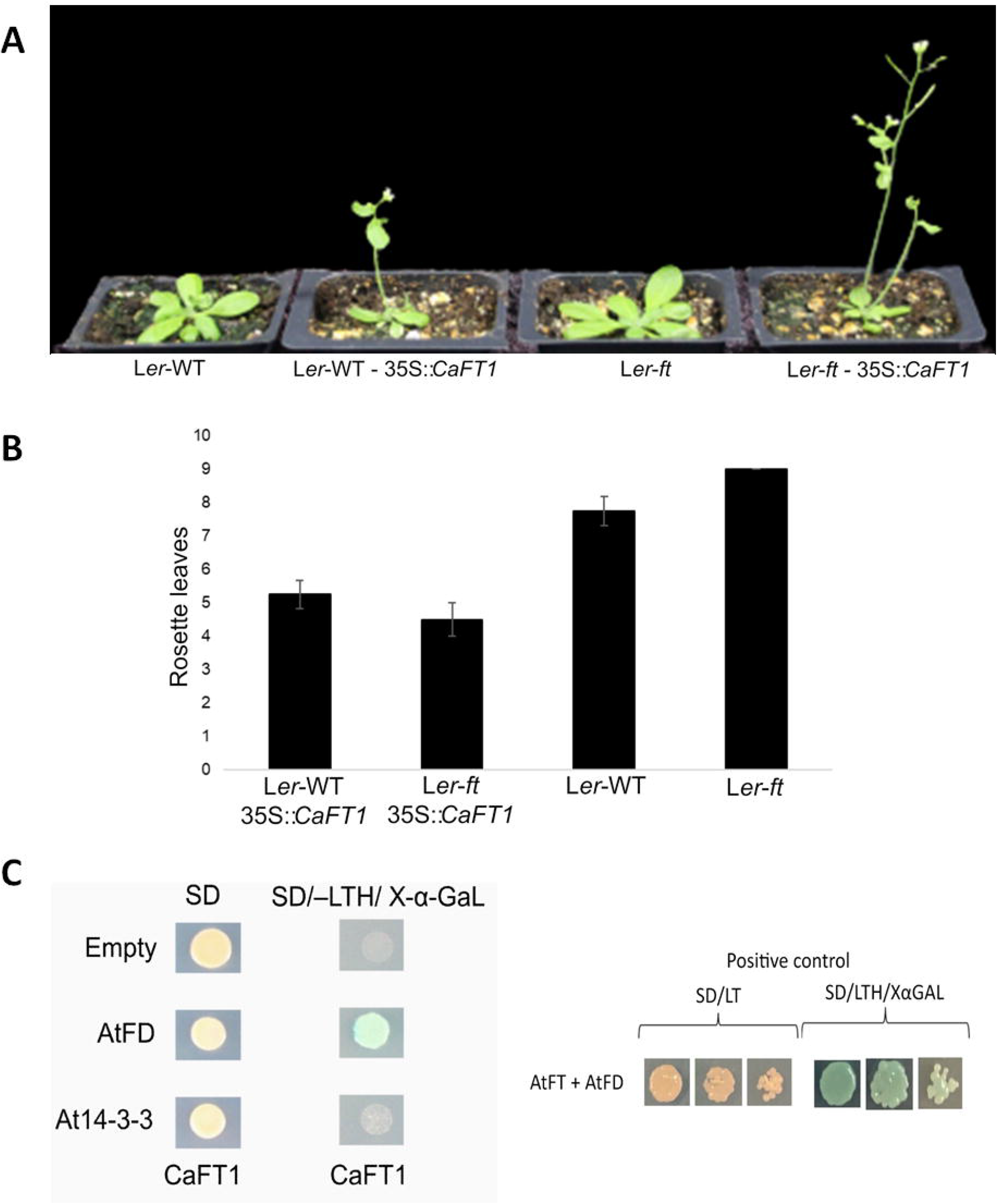
Functional analysis of *CaFT1*. A – Heterologous overexpression of *CaFT1* in *Arabidopsis thaliana* Ler ecotype causes early flowering. From left to right: Wild-type (L*er*-WT); 35S::*CaFT1* construct in L*er*-WT background; *FT* loss-of-function (L*er-ft*); 35S::*CaFT1* construction with L*er-ft* background. The overexpression of *CaFT1* driven by 35S CaMV was analyzed in 9 independent lines from 24 in T2 both L*er* wild type (L*er*-WT) plants as well as *ft* mutants. **B** – Rosette leaf number compared between L*er*-WT / 35S::*CaFT1,* L*er-ft* / 35S::*CaFT1,* L*er*-WT, and L*er-ft*. **C** - Yeast Two Hybrid Protein-protein interaction assay: Yeast transformed with Empty plasmid, *AtFD*, and *At14-3-3* inserted into pGADT7-RecAB plasmid (Prey), against *CaFT1* inserted into pBridge plasmid (Bait). Transformed yeast grown in SD medium as negative control and SD/-LTH/X-α-Gal as select medium and, as positive control Arabidopsis FT interacted with Arabidopsis FD in SD/-LTH/X-α-Gal select medium.

Known conserved components of the *Arabidopsis* Floral Activation Complex (FAC) include FT and FD together with a 14-3-3 protein (Kaneko-Suzuki *et al.*, 2018). Y2H assays with CaFT1 were carried out to test whether it interacts with AtFD and a 14-3-3 protein. CaFT1 protein was shown to interact with AtFD (**Fig. 1C**) as expected, since both proteins are reported as partners to form a protein complex (Abe *et al.*, 2005; Wigge *et al.*, 2005; Corbesier *et al.*, 2007; Jung *et al.*, 2016). However, despite the 14-3-3 protein being a component of the FAC with FT and FD proteins, no interaction was detected between CaFT1 and At14-3-3, suggesting diversification on 14-3-3 homologs and/or components of the FAC in *Coffea* sp. Together these results strongly support a role for CaFT1 as a floral inducer and that the selected *CaFT1* gene is a functional FT ortholog in *Coffea* sp.

### *CaFT1, CaFLC, CaCO, CaPIF4* show variable expression patterns over the course of a year

Cultivars of the same perennial species can show differential floral development patterns, especially flowering time (reviewed by Lopez *et al.*, 2021 in press). Such variation between cultivars is important for breeding programs to derive different cultivar genotypes with contrasting floral development times (Carvalho, 2008). Relevant to this study, variable flowering patterns were observed for the *C. arabica* cvs. Iapar 59, Acauã and *C. canephora* cv. Conilon cultivars (**Fig. S5**). To determine the floral induction window and explore a possible intra- and inter-specific diversification of *CaFT1* related to the contrasting flowering patterns of coffee cultivars, the expression of *CaFT1* in leaves of these genotypes were examined by RT-qPCR over the course of a year (**Fig. 2**).

**Figure 2:**
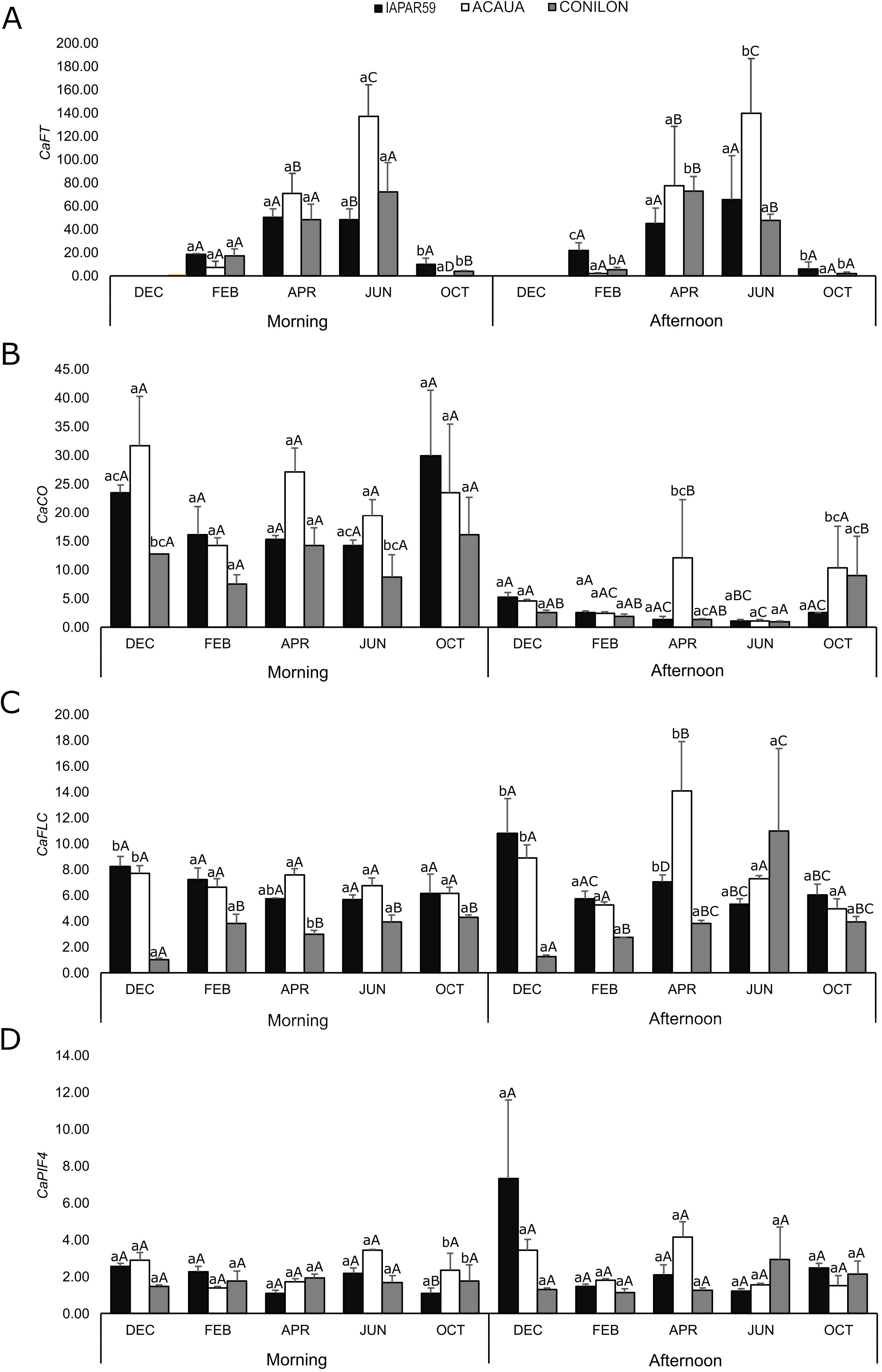
RT-qPCR expression analysis of putative floral regulators genes in leaves collected over one year from three different coffee genotypes: *C. arabica* cvs. Iapar 59 and Acauã and *C. canephora* cv. Conilon. Expression analysis were performed for the genes previously characterized (Fig. 1 and S1 to S4), *CaFT1* (A), CaCO (B), *CaFLC* (C) and *CaPIF4* (D). Leaf samples of different coffee genotypes were used. *C. arabica* cvs. IAPAR59 (black bars) and Acauã (white bars) and *C. canephora* cv. Conilon (gray bars). Samples collected at the first hours of the day (morning) and at the last hours of the day (Afternoon) in December of 2016 (DEC) and February (FEB), April (APR), June (JUN) and October (OCT) of 2017. Relative expression differences were calculated as log2 fold Change by Linear Mix Model as described by Steibel, *et al.* (2009). Letters at the top of the bars indicate the statistical differences, in which lower case letters represent comparisons between genotypes in each month, and capital letters compare the same genotype at different time points during the year.

Despite quantitative differences, all coffee genotypes showed a clear “bell shaped” pattern of *CaFT1* expression during the year for both morning and afternoon samples. *CaFT1* expression was observed in leaf samples from February, and increased progressively to maximum expression in June, followed by drastically reduced transcript levels in October and December (**Fig. 2A**). This result was correlated with the precise floral induction window, extending and encompassing the described emergence of floral buds (Majerowicz and Söndahl, 2005; de Oliveira *et al.*, 2014). Comparing genotypes, *C. arabica* cv. Iapar 59 and *C. canephora* showed similar values for *CaFT1* expression throughout the year and independent of the time period, whereas *C. arabica* cv. Acauã showed lowest expression levels in February and highest in April and June (**Fig. 2A**). This result can be associated with phenotypic differences between early- and late flowering coffee cultivars (**Fig. S5**).

Moreover, *FT* and florigen genes from diverse plant species have been shown to act as a hub for floral meristem activation by integrating different interconnected pathways responsive to environmental signals (Amasino, 2010; Andrés and Coupland, 2012). Thus, to gain insights into the regulation of *CaFT1,* the co-expression expression of photoperiod and thermosensitive FT regulators, as *CaCO*, *CaFLC* and *CaPIF4* (**Figs. S1 to S4)** were also determined at the same time points (**Fig. 2B, 2C and 2D**). Expression analysis showed that *CaFT1* transcription is detected first in leaves collected in February and increases gradually until it reaches a maximum in June. This period coincides with the shorter day length photoperiod and colder temperatures, typical of the winter in coffee-growing regions of Brazil (**Fig. 4A and S6**).

For all coffee genotypes, expression of *CaCO* was higher in the morning compared to afternoon (**Fig. 2B**), in agreement with the photoperiod-dependent regulation of *CO* orthologs reported in other plants (Suárez-López *et al.*, 2001; Zuo *et al.*, 2011). In the morning, *CaCO* expression presents a “smile shape” (or inverted bell shape) pattern throughout the year and was quite similar among genotypes, which means higher values in December and October and lower in the middle timepoints. This result correlates with the photoperiod of longer day length (**Fig. 4A and S6-A**). In afternoon samples, *CaCO* expression was lower and stable with few differences observed for genotypes, except in April and October. Comparing genotypes, in the morning *CaCO* was expressed at lower levels in cv. Conilon than *C. arabica* both cvs. for all time points throughout the year.

Regarding thermosensitive FT regulators, expression of *CaFLC* and *CaPIF4* was detected in all months for all three coffee genotypes, with few differences during the year in the morning, but varying in the afternoon (**Fig. 2C and 2D**). In the afternoon, *CaFLC* expression presents different patterns and maximum levels depending on genotype, for *C. arabica* cv. Iapar 59 it is in December and cv. Acauã in April and *C. canephora* cv. Conilon June. In the morning, no significant differences were found for *CaFLC* expression between *C. arabica* genotypes while cv. Conilon were always expressed less. Similar to *CaFLC*, *CaPIF4* expression in leaves showed little variation throughout the year, and no expression differences between coffee genotypes were observed during the day or throughout the year, except for cv. Acauã in October (**Fig. 2D**).

The *CaFLC* and *CaPIF4* expression did not vary with the temperature changes in Brazil (**Fig. 4A and S6-B), with** the lowest values in June and highest in October and December, except for *CaFLC* in June and afternoon for cv. Conilon. Moreover, and importantly, *CaFLC* and *CaPIF4* expression did not correlate with the expression pattern of *CaFT1* since it is expressed at higher levels in June when the putative negative regulator *CaFLC* is expressed and *CaPIF4* did not change. In addition, no differences or expression correlations were found for precipitation and humidity (**Fig. S6-C and D).** Thus, these results suggest that the floral inductive *CaFT1* pathway is not influenced, or at least to a lesser extent, by these thermosensitive floral FT regulators.

### Analysis of carbohydrate content in coffee leaves shows correlation with *CaFT1* expression

Sugars provide energy to all metabolic processes in plants and, with respect to the preparation for reproductive development, possibly expression of florigens such as *FT* as well (Corbesier *et al.*, 1998; Ohto *et al.*, 2001; Cho *et al.*, 2018). To examine a possible correlation with the *CaFT1* expression pattern and the energy status of coffee leaves, the carbohydrate content (total soluble sugar, sucrose, reducing sugar and starch) in adult plants was quantified at two daily timepoints over the course of a year.

Sucrose, starch, total soluble sugar and reducing sugars were quantified from the same samples used for expression analysis (**Fig. 2).** Results show patterns of distribution for the different coffee genotypes (**Fig. 3**). In general, all genotypes showed higher values for Total Soluble sugars (TS) and Sucrose (SC) in December and April (**Figs. 3A and B**), the summer period in Brazil, which is hotter and has more hours of daylight (**Figs. 4A**, **S6-A and B**). Similar results were observed for sucrose, but it was noted that cv. Iapar 59 had higher levels in all months (**Fig. 3B**). For reducing sugars (Fig. 3C), all genotypes had similar levels throughout the year, with *C. arabica* cv. Acauã showing higher values at all timepoints, except April. Finally, the quantification of starch levels showed a contrasting pattern between coffee genotypes. For example, the cv. Iapar 59 and cv. Conilon showed a clear pattern of increasing values from December to October in the next year, differing only at the highest timepoints, June and April, respectively. Whereas the cv. Acauã showed lower starch levels overall compared to other genotypes and the highest levels in October (**Fig. 3D**).

**Figure 3:**
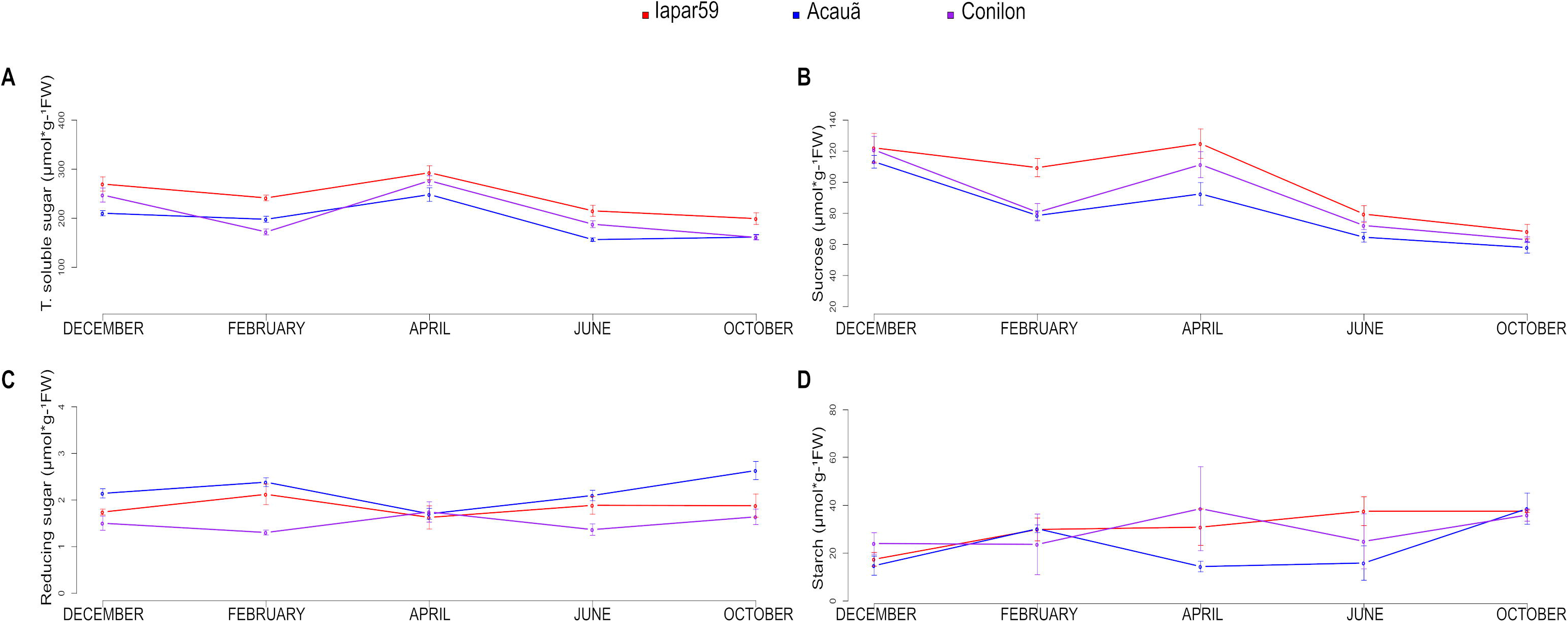
Carbohydrate content quantification from coffee leaves in microgram per gram of fresh weight. Samples collected from two *Coffea arabica* cultivars, Iapar 59 (red line), Acauã (blue line), and one *Coffea canephora* cultivar, Conilon (purple line), collected along five months of a year (DECEMBER, FEBRUARY, APRIL, JUNE, and OCTOBER). **A** – Total soluble sugar; **B** – Sucrose; **C** – Reducing sugar; **D** – Starch. Each sampling data set is represented by a point in the line with the standard deviation. All cultivars in each timepoint were represented by 3 biological samples and two technical replications.

**Figure 4:**
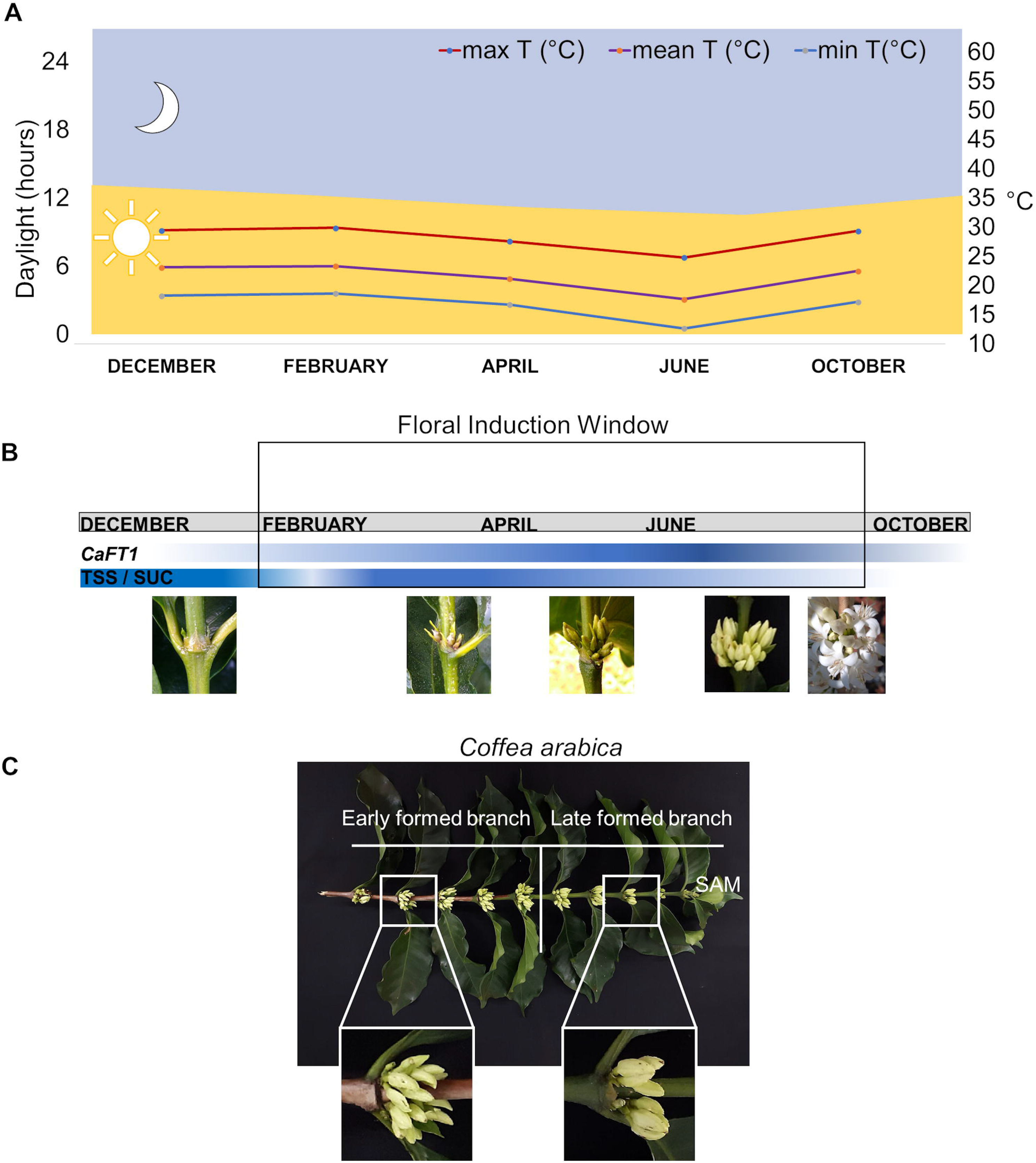
The floral induction window in coffee trees as a model for perennial species. Figure **A**, left axis of the graph represents the daylight hours variation divided by 24 hours which is separated in yellow color (day) and light blue color (night) in the graphic area. Right axis represents degree values (Celsius) from minimum (blue line), mean (purple line), and max (red line) temperature variation over the year; **B**, proposed floral induction window from February until October based on variation of *CaFT1* expression levels, total soluble sugars (TSS) and sucrose (SUC) content over the year. Gradients of color intensity (dark blue indicates higher level, light blue lower levels and white not detectable) represents the content level in terms of gene expression, total soluble sugar, and sucrose variation. The floral development stages observed in plants when the leaves were collected for the five time points (months) are also indicated in the figure; **C**, shows a plagiotropic branch with floral buds in the same development in both parts, the early formed branch (brown and lignified stem) and the late formed branch (green stem). Because the late formed branch originates from early nodes, the continuous vegetative activity of the shoot apical meristem (SAM) along the year (details in Figs. S5 e S6), in this figure suggests a need for extended florigen activity to induce axillary meristems formation at different times. Interestingly, floral meristems induced at different times reach flower buds at the same time.

These findings showed different patterns among the three coffee genotypes that could be related to differences observed for *CaFT1* expression (**Fig. 2A**) and, consequently, flowering time (**Fig. S5**). Thus, sucrose and other soluble sugars were the only types analysed whose levels correlated with the period of higher *CaFT1* expression, in April and June (Fig. 2A). Since sucrose is the main transport sugar (Lemoine, 2000) it is possible that *CaFT1* expression could be responsive to the sugar state. Similarly, a correlation between the energy status and flowering, as well as *FT* expression with trehalose-6-phosphate was established in *Arabidopsis* (Wahl *et al.*, 2013; Fichtner *et al.*, 2021).

## DISCUSSION

### *CaFT1* function and expression revealed the floral induction window in coffee that could be associated with the asynchronous flowering in perennial species

Previous studies focused on understanding the transition from juvenile to reproductive phase in perennial species have demonstrated that *FT*-related genes are involved in floral induction and dormancy time in response to seasonal stimuli in trees (Böhlenius *et al.*, 2006). However, the link between regulatory pathways of floral activation and asynchronous flowering is still poorly understood. First, in this work we demonstrate that overexpressing *CaFT1* in *Arabidopsis,* wild type and *ft-1* mutants, causes the typical early flowering phenotype and that CaFT1 interacts with AtFD (**Fig. 1**), a component of the floral activation complex (Kaneko-Suzuki *et al.*, 2018). Similar results were found for other annual or perennial plants when FT homologs were overexpressed in *Arabidopsis* and/or also in their respective species; for example, Poplar and apple (Kotoda *et al.*, 2010; Tränkner *et al.*, 2010), tomato (Cao *et al.*, 2016), *Eucalyptus* (Klocko *et al.*, 2016), Medicago (Laurie *et al.*, 2011) and blueberry (Gao *et al.*, 2016). Thus, CaFT1 can act as a florigen in *Arabidopsis* strongly suggesting same function in coffee plants as observed in other perennials. Further studies demonstrating that CaFT1 can act as a florigen in *Coffea* sp. will need to overcome the difficulty of transforming this species (Ribas *et al.*, 2011).

In coffee, the meristem floral transition window is reported to occur from January to March (Camargo and Camargo, 2001; Majerowicz and Söndahl, 2005). To understand the molecular events occurring at this transition, here we characterize an FT homolog from coffee that acts as a florigen and show that it is transcriptionally active for a longer period than the previously described window (Fig. 2A). *CaFT1* expression extends from February to October, reaching a maximum in June, which is a more precise characterization to indicate the potential floral induction window. Such a window overlaps with the entire period of floral bud development, which was interpreted as a continuum of induction (Fig. 4B). This extended window explains the formation of floral buds at the base of branches (older buds) together with induction of newly formed buds from the SAM at the distal end of branches (Fig. 4C). Floral bud development starts at different times until anthesis in September, when *CaFT1* expression decreases rapidly, followed by vegetative growth of new branches, restarting the cycle (**Fig. 4B and C**). Moreover, our finding suggests that the developmental time is also different, or, in other words, floral meristems induced early (at the base of the branch) develop slower than later ones (those closer to SAM). Thus, the *CaFT1* expression patterns observed in this study suggest that the prolonged expression window is associated with the asynchronous flowering behavior of coffee (**Fig. 4B and C**). Moreover, due to the conserved nature of flowering mechanisms, we speculate that this occurs in other tropical perennials plants.

In addition, the *CaFT1* expression pattern in coffee supports evidence that floral induction occurs only in axillary meristems because the FT from the leaves is distributed along the branch controlling floral induction to proximal meristems (McGarry and Ayre 2012). This also may explain the asynchronous flowering behavior of *Coffea sp* L., in which floral bud development occurs at axillary meristems from branches formed at different times and does not produce terminal flowers from the SAM (**Fig. 4C**). Alternatively, from the expression results (**Fig. 2 and 4B**), it is possible that *CaFT1* other functions in addition to floral induction, since it is reported to have a role in a wide range of developmental processes such as fruit set, vegetative growth, stomatal control and tuberization (Pin and Nilsson, 2012). Further study is required to explore this possibility.

### Coffee floral transition and photoperiodic stimulus

Despite the extended window of *CaFT1* expression, it is clear from the analysis with three genotypes that there is a conservation of the maximum level in June (Fig. 2), raising the question of whether there is a correlation with environmental cues during the period (Fig. 4). The florigen encoded by *FT* and its orthologs from numerous plants has been shown to induce the floral transition in response to temperature and photoperiod (Tsuji, 2017). With the isolation of an FT orthologue in coffee, this analysis can be extended to floral induction in tropical species that are subjected to high temperatures throughout the year. The expression profile of Ca*FT1* (**Fig. 2A**) may show less variation with respect to photoperiod effects, possibly because there is less daylight variation in equatorial regions compared to temperate zones.

Examination of flowering control in trees provides evidence that CONSTANS (CO) protein is responsible for *FT* ortholog expression induction by its accumulation in long days (LD) (Böhlenius *et al.*, 2006). However, despite *CaCO* showing expression variation in response to day length (**Fig 2B**), its expression over the course of the year showed an inverse relationship. The *CaCO* showed a variable expression pattern for all three coffee genotypes with higher values reached in the morning, according to the circadian clock that is mainly responsive to photoperiod (Valverde *et al.*, 2004). Comparing expression over the year, *CaCO* was higher in December and October (**Fig. 2B**), coinciding with summer in the Southern hemisphere and the period of greatest solar light incidence **(Fig. 4 and S6-A)**. This was opposite to the pattern since *CaFT1* expression is higher during the Brazilian winter (Fig. 2A), with the shortest day length (**Fig. 4A and S6-A**) and more highly expressed in summer in December and October (**Fig. 2A**). Other factors might be involved in the CO regulatory pathway of *CaFT1* expression; for example, Wenkel, *et al.*, (2006) showed that the protein HEME ACTIVATOR PROTEIN2 (HAP2) or HAP3 can create a complex with CO that causes a reduction in the expression of *FT*.

### Expression of *CaFT1* is not correlated with *CaFLC* and *CaPIF4* expression, suggesting alternative thermoregulatory pathways that are not influenced by temperature

In regions where temperatures vary during the year, floral initiation and dormancy for some species can be controlled by cold exposure or vernalization (Michaels *et al.*, 2005; Kim *et al.*, 2009; Madrid *et al.*, 2021). In the *Arabidopsis* vernalization pathway, *FLC* expression is repressed after prolonged exposure to colder temperatures, thus activating *FT* expression and allowing floral meristem induction (Michaels and Amasino, 1999; Helliwell *et al.*, 2006; Aikawa *et al.*, 2010). On the other hand, *PIF4* is responsive to warmer temperatures and positively regulates *FT* expression (Kumar *et al.*, 2012). Thus, we hypothesized that FLC and PIF4 homologs in coffee could be related to thermal-sensitive pathways, possibly regulating *CaFT1* expression. Here we use them as a probe to evaluate co-expression patterns that are integrated with temperature changes during the growing season.

In this work, *FLC* homologs were found in coffee, which is interesting given that it is a tropical plant that does not normally experience freezing temperatures. In line with this, *CaFLC* expression was not associated with temperature variation, as might be expected. A similar result was found for *CaPIF4* expression (Fig. 2C and D, Fig. 4A and S6-A). Over the course of the experiment, when the lowest temperatures were registered in June and the highest occurred in December, *CaFLC* and *CaPIF4* showed stable expression patterns or changes in different periods not correlated with these temperature changes. Moreover, comparing *CaFT1* and *CaFLC* expression patterns, there was no inverse co-expression correlation as would be expected if FLC repressed *FT* activity, nor was there a positive correlation with *CaPIF4,* suggesting it is not an FT inducer. Overall, expression of *CaFT1* was highest in June, during the Brazilian winter (Fig. 2A, Fig. 4A and S6-A), suggesting positive regulation by cold or even drought, typical characteristics of the Brazilian winter. Further analysis under controlled conditions will demonstrate whether *CaFT1* is responsive to cold and/or drought. In addition, *CaFLC* expression varied between coffee genotypes, with Iapar 59 and Acauã showing similar expression, whereas Conilon showed lower expression levels throughout the growth period and higher in afternoon samples of June (Fig. 2C). These differential expression patterns suggest intra- and inter-specific transcriptional differences, which might coincide with contrasting flowering behavior (DaMatta and Ramalho, 2006), or it could be associated with greater phenotypic homeostasis of the allotetraploid *C. arabica* than its diploid parents in response to different temperature conditions (Bertrand *et al.*, 2015).

These results show no evidence that the *FT-*dependent floral transition pathway is regulated by coffee *FLC* or *PIF4* homologs, thus the potential roles of *CaFLC* and *CaPIF4* in flowering of tropical perennials awaits further study. Alternatively, despite tropical species such *Coffea* sp. having sequences that are homologous to Arabidopsis *FLC* and *PIF4,* they may have functions not necessarily related to floral control in response to temperature variation. The finding that *CaFLC* expression varied widely in different coffee tissues such as, roots, leaf, SAM at all floral and fruit developmental stages (de Oliveira *et al.*, 2014) suggests functions other than flowering repression, such as coordinating organogenesis together with SOC1 as reported in *Arabidopsis* (Deng *et al.*, 2011). In support of a possible role in organogenesis, *CaFLC* expression is upregulated in response to drought (Barreto *et al.*, 2012), which coincides with the growth latency in the G4 stage interpreted as a dormant stage (Wormer and Gituanja 1970; Majerowicz and Söndahl, 2005). At present, however, no direct mechanism for this has been described yet (reviewed by Lopez *et al.*, 2021 in press).

Future research will decipher whether CaFLC and CaPIF4 are involved in flowering, and their relationship with environmental signals and the FT-regulatory pathways. In addition, both genes are regulated by chromatin epigenetic changes in Arabidopsis (Helliwell *et al.*, 2006; Kumar *et al.*, 2012; Madrid *et al.*, 2021), a very little explored field of research in *Coffea* sp. and crop perennials in general.

### The role of carbohydrates in coffee floral induction as a model for perennials

Sugars have been shown to be important chemical signals that affect flowering, as strongly supported in model plants like *Arabidopsis* (Wahl *et al.*, 2013; Cho *et al.*, 2018; Fichtner *et al.*, 2021). Since no correlation was found between *CaFT1* expression and possible environment-related regulators, we examined sugar levels to assess their role as possible regulators in tropical perennial species. Gene expression analysis showed association with Total Soluble Sugar and Sucrose levels (**Fig. 3A and B**) in relation to temperatures during the five analyzed months, showing an association with warmer periods of the year (**Fig. 4A and S6-A**). This association is related to higher levels of TS and SC and coincided with *CaFT1* expression levels (**Fig. 2A**). As previously described by Cho *et al.* (2018) and, in accordance with our results (**Fig. 4**), sugar levels, a product of photosynthesis, can change according to seasons in response to more hours of light and higher temperature.

Sugar variation is an important indicator of reproductive phase initiation, with evidence that it affects the expression of floral integrators, such as *FT* (Rolland *et al.*, 2006; Moghaddam and Van den Ende, 2013). In our study, sugar accumulation was higher in December and April, the summer period in Brazil that is hotter and has more hours of daylight (**Fig. S6-A**). Interestingly, these results coincide with two relevant periods associated with *CaFT1* expression - in December, before the increase in *CaFT1* expression in February, and in April which is a period before *CaFT1* expression peaks in June. Whether this pattern suggests that sugars act as a stimulus for *CaFT1* expression requires further evidence (**Fig. 1A**). Association of *CaFT1* with TS and sucrose levels suggest that its expression is responsive to carbohydrate signals, showing a possible connection between sugars and floral induction in perennial species. Thus, it will be interesting to determine whether energy status is a key regulator of *CaFT1* expression and controls coffee floral development, further suggesting that environmental factors are indirectly involved since photo-assimilate production is affected by photoperiod and temperature (Pego *et al.*, 2000; Lastdrager *et al.*, 2014). Accordingly, we demonstrate that sugar content in coffee plants changes in response to temperature regimens (de Oliveira *et al.*, 2020). These findings support previous studies demonstrating a correlation between *FT* expression pattern and carbohydrate content (Corbesier *et al.*, 1998).

## CONCLUSIONS

Floral induction in perennial plants is a poorly understood process that integrates endogenous and environmental factors. Elucidating the molecular components of this floral regulatory process is a necessity in the face of imminent climate change to secure food production (Howden *et al.*, 2007; Lobell *et al.*, 2011; Zhao *et al.*, 2017). In this work we identified the coffee FT ortholog and determined its expression profile to describe the precise and potential floral induction window. This analysis includes the analysis of key *FT* regulators responsive to environmental signals, evaluation of climatic parameters and sugar content over the period of one year. Together, our results indicate a continuum of florigen transcription, conserved between contrasting coffee genotypes, that could underlie asynchronous floral development and flowering. The environment-related floral regulators, *CaCO*, *CaFLC* and *CaPIF4,* were not co-expressed as expected with *CaFT1*, whereas a correlation was found with sugar content, which is affected by environmental changes over the course of a year. This suggests that *CaFT1* is not directly regulated by these genes, but that there may be a connection between sugar metabolism and florigen function in coffee. Thus, the present work contributes to comprehending asynchronous flowering in tropical perennials plants and provides a basis for targeting molecular components in crop breeding programs.

## Abbreviations

BFT: BROTHER OF FT
CO: CONSTANS
FLC: FLOWERING LOCUS C
FM: Floral meristem
FT: FLOWERING LOCUS T
GA: Gibberellin
LD: Long day
MFT: MOTHER OF FT
PEBP: Phosphatidylethanolamine-binding protein
PIF4: PHYTOCHROME INTERACTING FACTOR 4
SAM: Shoot apical meristem
SFT: TWEEN SISTER OF FT
SD: Short day
TFL1: TERMINAL FLOWER 1
VM: Vegetative meristem

## SUPPLEMENTARY DATA

Table S1 shows all the primers used in this work; figures S1 to S4 show the phylogenetic analyses and amino acids alignments for the gene families *FT*, *CONSTANS*, *FLC* and *PIF4*; Figure S5 shows a photo panel of the plagiotropic branches with floral meristems at different stage of development comparing *C. arabica* cvs. Iapar 59 and Acauã and *C. canephora* cv. Conilon in three different timepoints; Figure S6 shows the variation of photoperiod, temperature, precipitation and relative humidity during the experiments at the experimental field of Federal University of Lavras (UFLA, MG/Brazil).

## ACKNOWLEDGEMENTS

The authors thank the members of the Laboratory of Plant Molecular Physiology (LFMP, UFLA/Brazil) for structural support of the experiments; Prof. Dr. Mario Lucio (UFLA/Brazil) and Instituto Brasileiro de Ciência e Tecnologia do *Café* (INCT-Café) to support plant material and field experiments; Fundação de Amparo à Pesquisa do Estado de Minas Gerais (FAPEMIG, grant number CAP APQ 03605/17), Conselho Nacional de Desenvolvimento Científico e Tecnológico and INCT-Café for the financial support; Prof. Dr. Paulo Eduardo R. Marchiori and colleagues from his lab (UFLA/Brazil) for the support on sugars and proteins analyses. Research in JCs lab is supported by a Natural Sciences and Engineering Research Council Discovery grant. Mike Mucci and Leane Illman from the Phytotron facility at the University of Guelph provided expert plant care.

## AUTHORS’ CONTRIBUTIONS

C.H.C., R.R.O. and A.C-J. conceptualized the project. C.H.C., conducted all experiments and data analyses. L.P.P. participated in the collection of plant materials, RNA extractions and the RT-qPCR execution. V.L. and C.F. participated in the yeast two hybrid and heterologous expression analyses. T.H.C.R. assisted bioinformatic and statistical analyses. R.R.O., J.C. and A.C-J. conceived of and supervised the experiments and data analyses. C.H.C. wrote the manuscript. R.R.O., J.C and A.C-J revised the manuscript and contributed to writing.

## DATA AVALIABILTY

Main data supporting the findings of this study are available within the paper and within its supplementary materials published online. The raw data used for analyses and figures are available from the corresponding author, Antonio Chalfun-Júnior, upon request.

