## Supplemental data for "Expression of coffee florigen *CaFT1* reveals a sustained floral induction window associated with asynchronous flowering in tropical perennials"

**SUPPLEMENTARY MATERIAL**

**Tables**

**Table S1 –** Primers used for gene isolation, plant transformation and RT-qPCR analyses.

| Objective | Gene | Forward 5’ – 3’ | Reverse 5’ – 3’ |
| --- | --- | --- | --- |
| PCR/Cloning | *CaFT1* | ATGCCTAGAGGGGGAGGAGA | TTATCGTCTTCTGCCTC |
| RT-qPCR | *CaFT1* | GTTCACCAGGTCCCTAAGCC | ATCGTCACCTCCAATCTCAACC |
| RT-qPCR | *CaCO* | TGCTATTTGGTGGCGAGGTT | CGCTGTCTCCTCCGTAACTC |
| RT-qPCR | *CaFLC* | GACGGGTCTGCAAGAATTAGC | GTGGTCAACATCTGGCTCCT |
| RT-qPCR | CaPIF4 | CACTACATGCCCCGCTTAG | GAACATTGTGGTTTGGTGGCA |

**Figures**

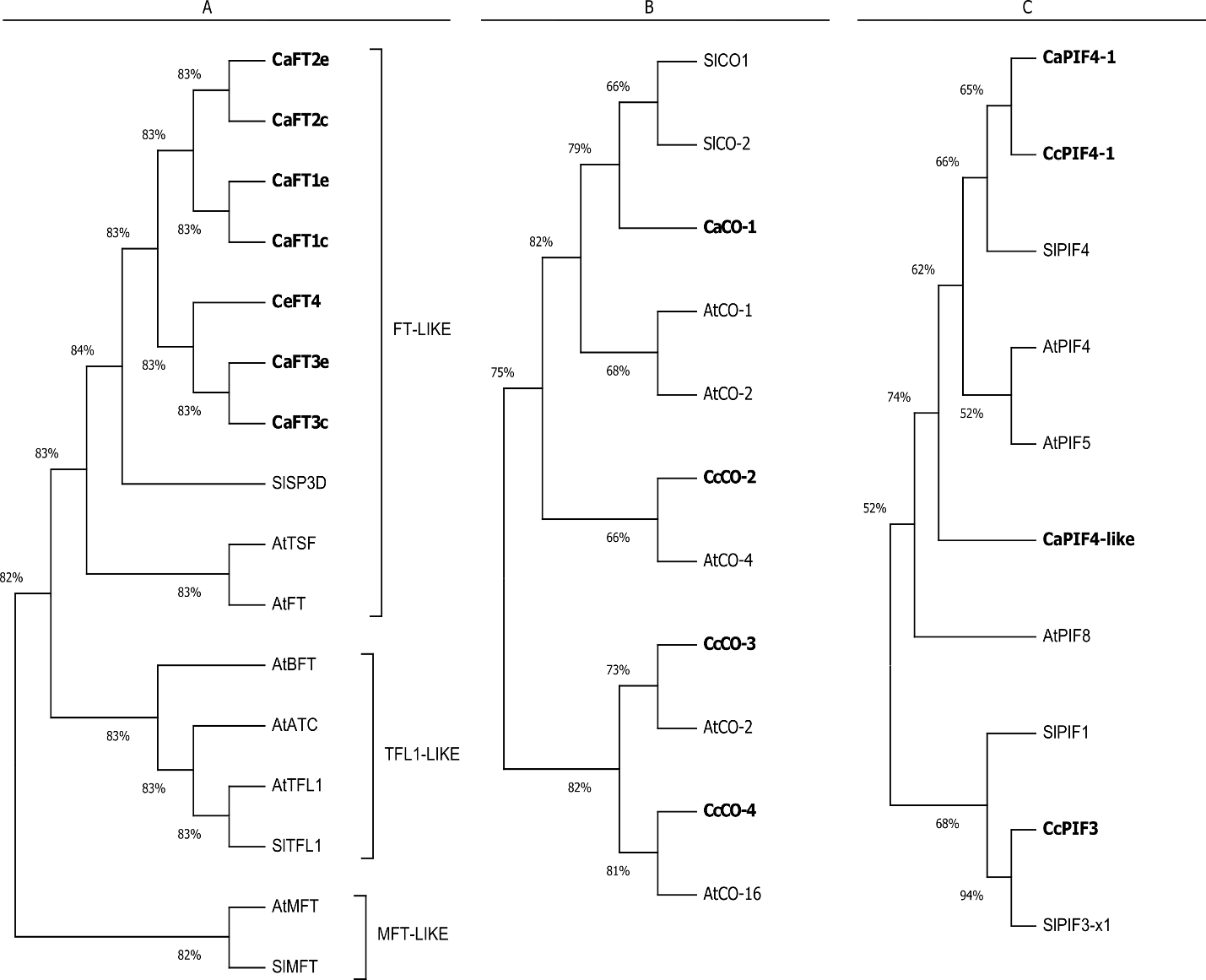

**Figure S1 –** Phylogenetic trees relating protein sequences of *Coffea sp*. to different homologs in *Arabidopsis thaliana* (At) and *Solanum lycopersicum* (Sl). Tree showing the main subgroups reported for the *FT* family, FT-like, TFL1-like and MFT-like; B) Tree for CONSTANS (CO) homologs; C) Tree for PHYTOCHROME INTERACTING FACTOR (PIF4) homologs. Coffee sequences are identified in bold and were named according the following criteria: i) genome species dataset, *C. arabica* (*Ca*), *C. canephora* (*Cc*) and *C.* [*eugenioides*](https://www.google.com/search?client=ubuntu&hs=lSC&channel=fs&q=C.+eugenioides&spell=1&sa=X&ved=2ahUKEwjrzpSEpLruAhUYJbkGHaioD6AQBSgAegQIBxA1) (*Ce*); ii) name of the closest homolog; iii) number of copies found in the genome (i.e. CaFT1 to 4); iv) adding a letter “c” or “e” to indicate the *C. arabica* sub-genome copy, *C. canephora* and *C.* [*eugenioides*](https://www.google.com/search?client=ubuntu&hs=lSC&channel=fs&q=C.+eugenioides&spell=1&sa=X&ved=2ahUKEwjrzpSEpLruAhUYJbkGHaioD6AQBSgAegQIBxA1)*,* respectively*.* Thus, the selected sequences were (gene ID in parenthesis): CaFT1c (XP_027087615.1), CaFT2e (XP_027090020.1), CaFT2c (XP_027093486.1), CaFT3e (XP_027083083.1), CaFT4e (XP_027079832.1), CaCO-1 (XP_027076653.1), CcCO-2 (Cc04_g07160), CcCO-3 (Cc05_g05160), CcCO-4 (XP_027084679.1), CcPIF4-1 (Cc05_g00300) and CaPIF4-1 (XM_027205516.1).

**Figure S2 –** Alignment of the predicted amino acid sequences belonging to the CETS family from *Coffea arabica* (*Ca*), *Coffea eugenioides* (*Ce*), *Arabidopsis thaliana* (*At*) and *Solanum lycopersicum* (*Sl*). The figure shows conserved amino acids and motifs amongst sequences of the same reported subgroup, FT-like, TFL1-like and MFT-like (Karlgren et al., 2011; Nasim et al., 2017), which supports duplication events observed in the figure S1. Sequences were aligned using the Genedoc tool (www.psc.edu/biomed/genedoc) in which the grade of amino acids conservation is represented by a color scale, being black boxes the highest degree and no color the lowest. A conserved amino acid considered necessary for the floral activation function of FT-like genes, Y85 (Hanzawa et al., 2005), is identified by a red box in the figure.
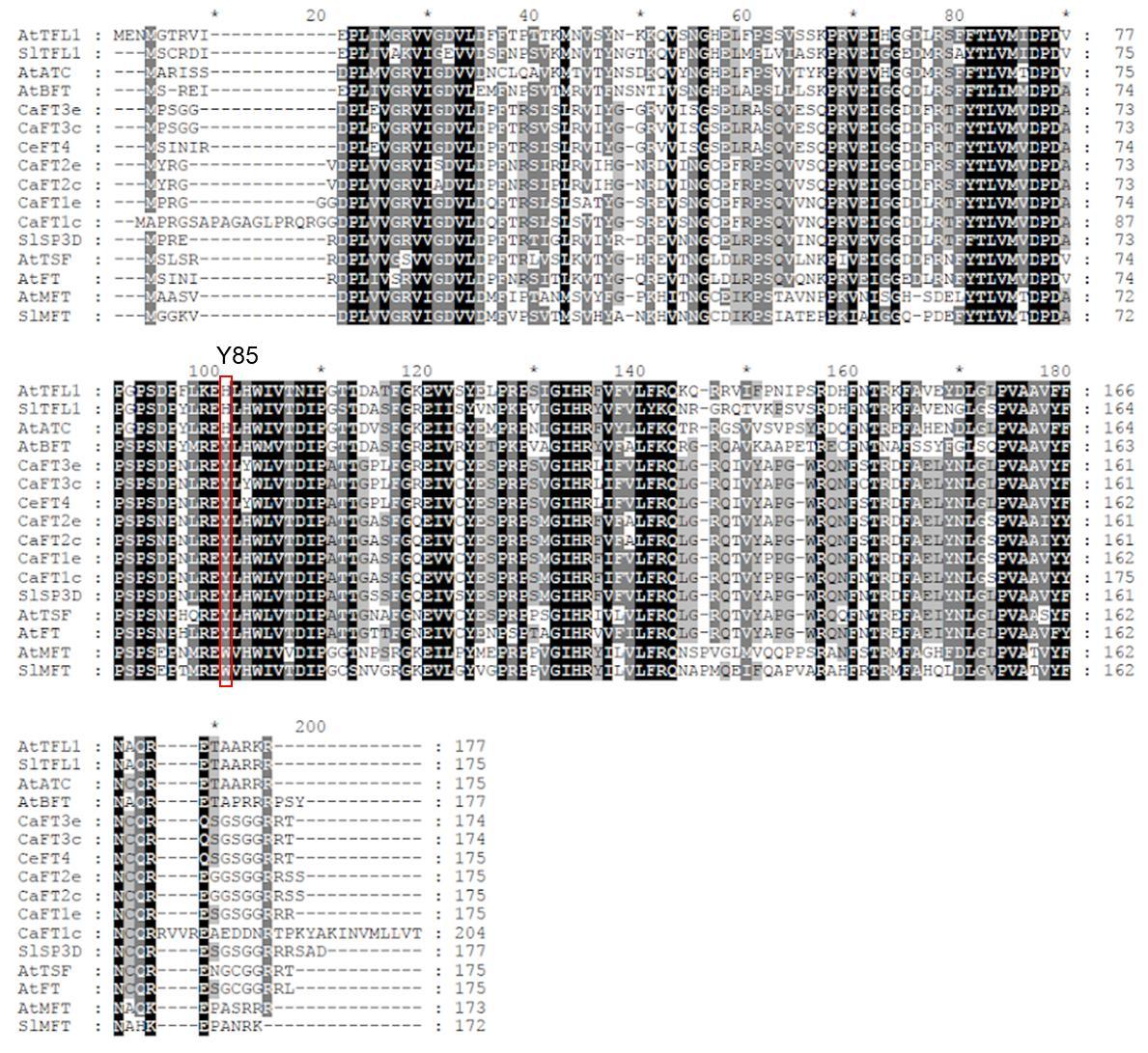

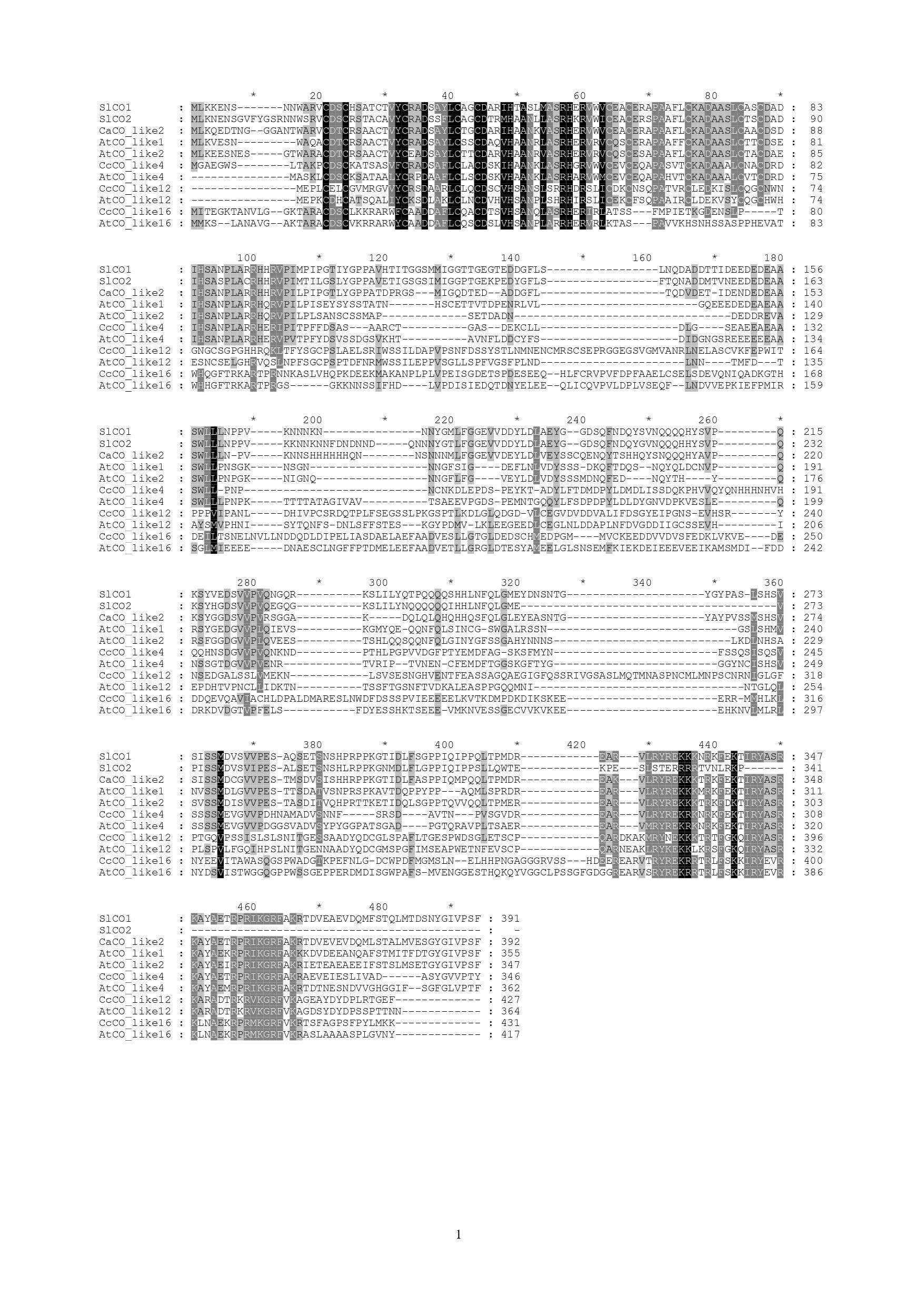

**Figure S3** – Alignments of predicted amino acid sequences belonging to the *CONSTANS* gene family from *Coffea arabica* (*Ca*), *Coffea eugenioides* (*Ce*), *Arabidopsis thaliana* (*At*) and *Solanum lycopersicum* (*Sl*). Sequences were aligned by using the Genedoc tool (www.psc.edu/biomed/genedoc) showing the grade of amino acids conservation. Shared sequence similarity is represented by a color scale intensity (black boxes represent the highest degree of similarity and no color the lowest).

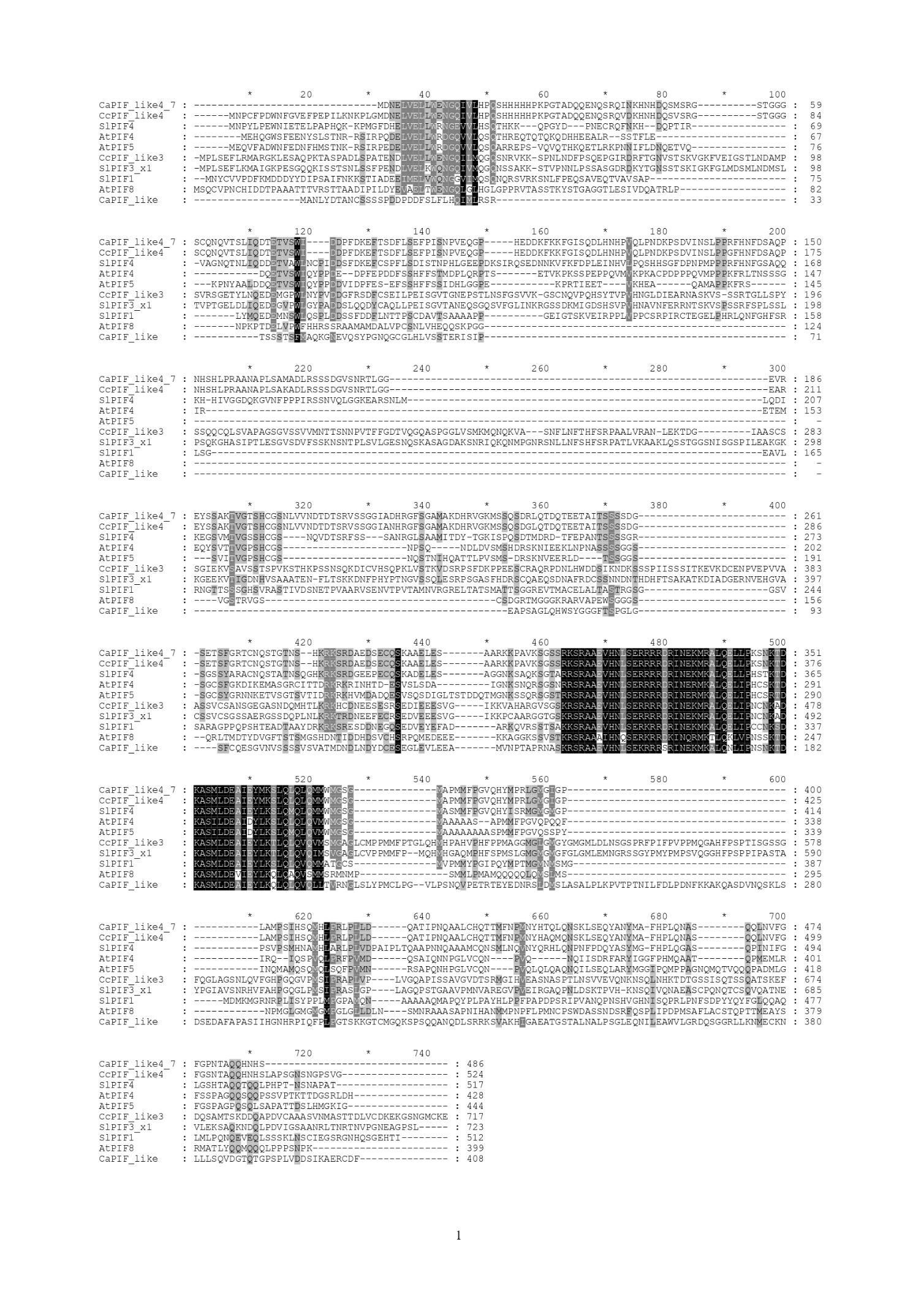

**Figure S4 –** Alignments of predicted amino acid sequences belonging to the *PHYTOCHROME INTERACTING FACTOR* 4 (*PIF4*) gene family from *Coffea arabica* (*Ca*), *Coffea eugenioides* (*Ce*), *Arabidopsis thaliana* (*At*) and *Solanum lycopersicum* (Sl). Sequences were aligned by using the Genedoc tool (www.psc.edu/biomed/genedoc) showing the grade of amino acids conservation. Shared sequence similarity is represented by a color scale intensity (black boxes represent the highest degree of similarity and no color the lowest).

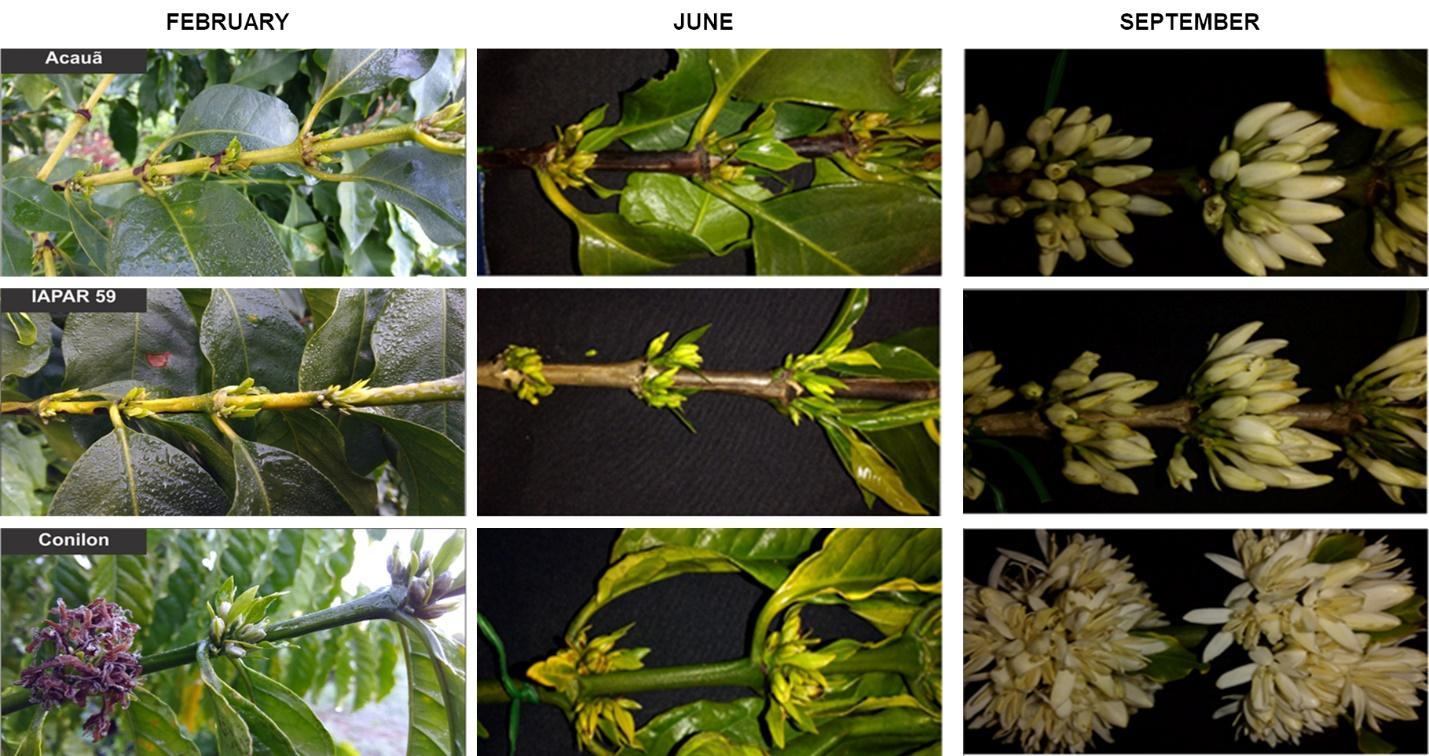

**Figure S5** **–** Orthotropic branches of *C. arabica* cvs. Iapar 59 and Acauã and *C. canephora* cv. Conilon, observed in February, June, and September. The cv. Conilon presents an accelerated floral development in relation to the cvs. Iapar 59 and Acauã.

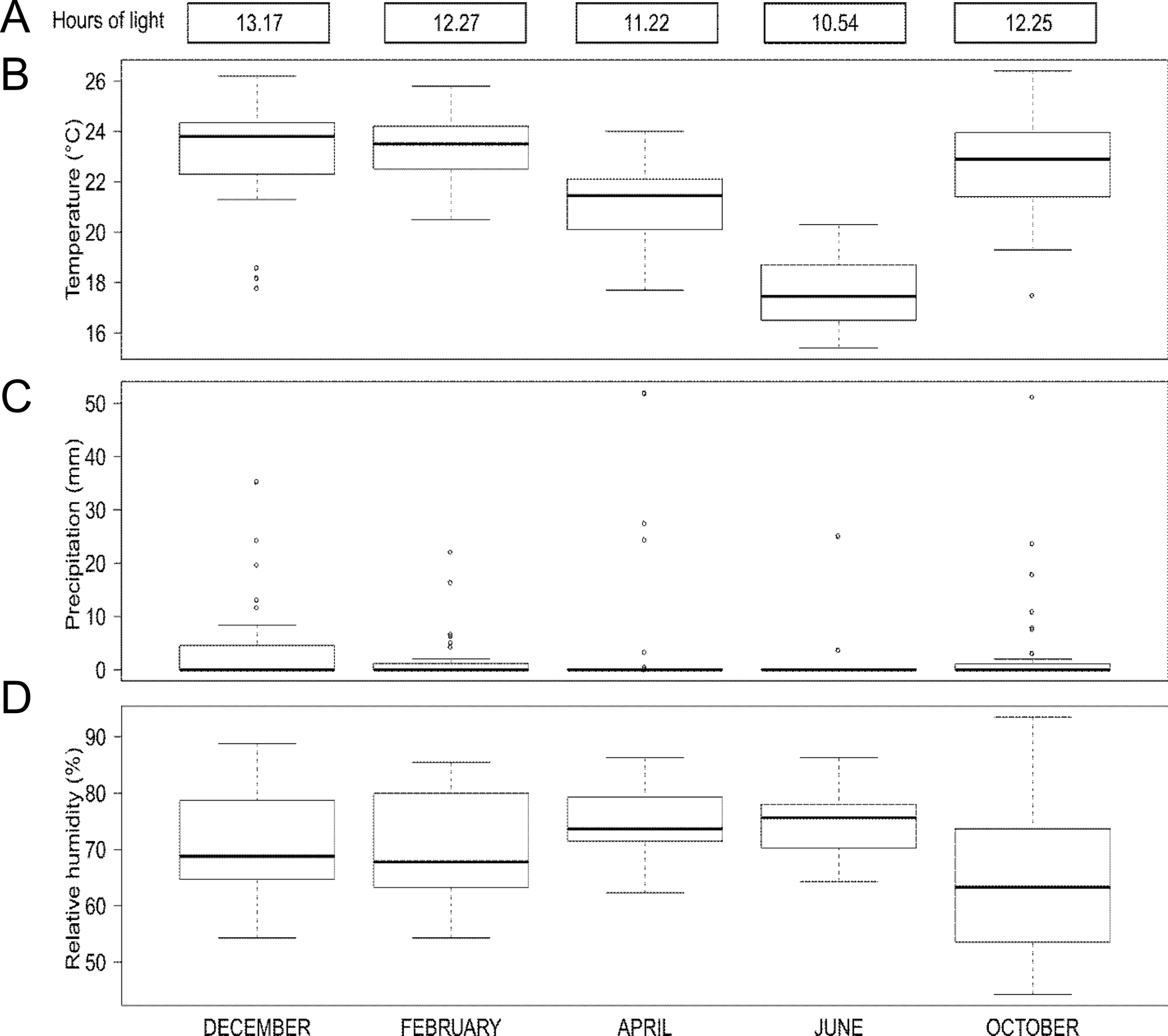

**Figure S6** – Variation of environmental conditions in five points of the year at the experimental field of Federal University of Lavras (UFLA, MG/Brazil). **A** – Top figure shows the daylight hours when the leaf samples were collected each month; B) Box plot representation of Temperature (°C) mean variation C) Box plot representation of the Precipitation (mm) mean variation; D) box plot representation of Relative humidity (%) mean All data was collected in December, February, April, June and October.
